# The modulatory effects of *Hericium erinaceus* and donepezil on amyloid accumulation and autophagy-associated clearance in Aβ_1-42_-induced N2a cells

**DOI:** 10.64898/2026.10.06.757182

**Authors:** Namatshego Moselakgomo, Wenzile Mthethwa, Sanet Van Zyl, Marieka Gryzenhout, Gabre Kemp, Claudia Ntsapi-Combrink

## Abstract

**Background and objective:** Alzheimer’s disease involves imbalanced amyloid-β (Aβ) production and clearance, supporting investigation of adjuncts to symptomatic treatment. This study evaluated whether methanolic *Hericium erinaceus* extract could support recovery after Aβ_1-42_-induced injury, alone and with donepezil, through modulation of β-secretase (BACE1) activity or autophagy-associated clearance.

**Methods:** Extract composition and BACE1 inhibition were assessed using liquid chromatography-mass spectrometry (LC-MS) and a fluorescence resonance energy transfer assay, respectively. Neuro-2a cells exposed to Aβ_1-42_ (2.42 µM) for 6 hours received individual or combined treatments for 24 hours. Cell viability was assessed using WST-1, combination interactions using Bliss independence and protein responses using Western blotting with paired bafilomycin A1 conditions across three independent experiments.

**Results:** LC-MS identified ergothioneine and trace hericenone D. The extract produced an apparent concentration-dependent BACE1-inhibitory response (IC_50_, 69.97 ± 36.10 µg/mL) and significantly increased post-injury viability at 40 µg/mL (150.2 ± 12.11% of the Aβ-induced control; *p* = 0.0070). Although the full combination matrix was predominantly additive (Bliss score, 1.95), 51.1 µM donepezil with 24.9 µg/mL extract produced local synergy (score, 34.29) and the highest observed viability. This pair showed the most coordinated directional protein pattern, with lower mean levels of amyloid precursor protein, Aβ and BACE1, and p62/Aβ accumulation following lysosomal blockade. Western blot differences were not statistically significant.

**Conclusion:** These integrated findings support *H. erinaceus* as a candidate adjunct to donepezil, with possible complementary effects on amyloidogenic processing and intracellular clearance. Further validation of autophagic flux is needed to confirm the proposed dual response.

## 1. INTRODUCTION

Alzheimer’s disease (AD) is a progressive neurodegenerative disorder and the most common cause of dementia. In 2021, an estimated 57 million people lived with dementia worldwide, with AD accounting for approximately 60-70% of cases (WHO, 2026). Amyloid-β (Aβ) accumulation is a central feature of AD and reflects an imbalance between production and clearance. Aβ is generated through sequential cleavage of amyloid precursor protein (APP) by β-site APP-cleaving enzyme 1 (BACE1) and γ-secretase (Vassar *et al*., 1999), while impaired clearance contributes to its accumulation in AD (Mawuenyega *et al*., 2010).

Macroautophagy, hereafter termed autophagy, is a lysosomal degradation pathway that supports intracellular protein and organelle turnover. This process is particularly important in long-lived neurons, which depend on sustained removal of damaged cellular components to maintain their function (Hara *et al*., 2006). Effective autophagic clearance requires cargo delivery by autophagosomes, followed by lysosomal fusion, acidification and degradation. Defects in these steps may therefore sustain intracellular protein accumulation despite ongoing autophagosome formation. In AD mouse models, defective autolysosomal acidification has been associated with neuronal accumulation of Aβ and APP-derived fragments before extracellular plaque deposition (Lee *et al*., 2022). These findings implicate impaired autophagic clearance in amyloid-associated injury and support investigating clearance alongside Aβ production.

The need to address these amyloid-related cellular abnormalities provides a rationale for investigating adjuncts to donepezil, a cholinesterase inhibitor used for symptomatic management of AD (WHO, 2026). *Hericium erinaceus*, an edible medicinal mushroom, is a candidate for such investigation because its constituents and preparations have shown neurotrophic and neuroprotective activities in experimental studies. Selected hericenones have stimulated nerve growth factor (NGF) synthesis *in vitro* (Kawagishi *et al*., 1991), while polysaccharide pretreatment attenuated Aβ-induced injury in PC12 cells (Cheng *et al*., 2016), and erinacine A-enriched mycelial preparations reduced amyloid plaque burden and increased insulin-degrading enzyme levels in APP/PS1 mice (Tzeng *et al*., 2016). These observations support evaluating *H. erinaceus* as a potential adjunct, although evidence from different preparations and protective studies does not establish its capacity to support rescue after established injury.

This study therefore investigated whether a methanolic *H. erinaceus* extract could support cellular rescue after Aβ_1-42_-induced injury, alone and alongside donepezil, and whether its effects could involve BACE1 inhibition, autophagy-associated clearance, or both. Extract composition, BACE1-inhibitory activity and basal cytotoxicity were assessed, followed by evaluation of single-agent rescue and combination interactions in murine Neuro-2a (N2a) cells. Amyloid-associated and autophagy-related proteins were examined to explore the cellular responses accompanying treatment.

## 2. MATERIALS AND METHODS

### 2.1 *H. erinaceus* extraction

Dried *H. erinaceus* powder was supplied by Prof. M. Gryzenhout, University of the Free State, South Africa. Extraction was adapted from Li *et al*. (2012): 1.012 g powder was stirred in 10 mL methanol for 24 hours at room temperature, filtered through Whatman No. 1 paper and rinsed with 10 mL methanol. Filtration continued for 24-48 hours. The filtrate (7.5 mL; 38.8 mg/mL) was stored at 4°C, diluted in dimethyl sulfoxide (DMSO) to 10 mg/mL and filter-sterilised (0.22 µm).

### 2.2 Donepezil preparation

Donepezil hydrochloride (≥98%; Sigma-Aldrich, D6821-50MG) was dissolved in anhydrous DMSO (20.8 mg/5.00 mL) to prepare a 10 mM stock, filter-sterilised (0.22 µm), aliquoted and stored at 4°C protected from light. Working solutions were prepared immediately before use.

### 2.3 Aβ_1-42_preparation

For model optimisation, freeze-dried Aβ_1-42_ (Sigma-Aldrich, AG974-500UG) was reconstituted at 1 mg/mL in deionised water/PBS (1:5, v/v), incubated at 37°C for 48 hours and diluted with anhydrous DMSO to 100 µM. Aliquots were stored at 4°C and diluted in complete culture medium immediately before use.

For downstream experiments, Aβ_1-42_ was dissolved in hexafluoroisopropanol (10 mg/mL), aliquoted and dried completely, following Nazere *et al*. (2022) with modifications. Peptide films were reconstituted in DMSO at approximately 2 mg/mL and diluted in complete culture medium to 2.42 µM.

### 2.4 LC-MS analysis

Targeted liquid chromatography-mass spectrometry (LC-MS) assessed ergothioneine and hericenone D using a SCIEX 4500 QTRAP coupled to a Shimadzu HPLC. Samples (10 µL) were separated on a Synergi MAX-RP C18 column (50 × 2.0 mm, 4 µm; Phenomenex) at 300 µL/min. Water and acetonitrile, each containing 0.1% formic acid, formed the mobile phases; the gradient lasted 3 minutes within a 10-minute run. Electrospray ionisation used a 500°C source temperature. Scyros 3.4.0 processed the data, with identification supported by retention times and reference standards. Untargeted profiling used positive- and negative-ionisation modes and spectral-library matching. Annotations remained tentative without confirmatory MS/MS.

### 2.5 BACE1 inhibition assay

The BACE1 FRET Assay Kit, Red (Thermo Fisher Scientific, P2985) was used following the manufacturer’s protocol and Mthethwa *et al*. (2026). Extract concentrations of 3.125-100 µg/mL were tested with BACE1 (1 U/mL working solution) for 90 minutes at room temperature. Fluorescence was measured at 520/585 nm excitation/emission using a GloMax Discover reader (Promega).

### 2.6 Cell culture

Wild-type mouse neuroblastoma cells (N2aWT; ATCC CCL-131) were cultured as previously described (Ntsapi and Loos, 2021; Mthethwa *et al*., 2026) in high-glucose Dulbecco’s Modified Eagle Medium containing 10% fetal bovine serum and 1% penicillin-streptomycin at 37°C and 5% CO_2_. Cells were passaged twice weekly using 0.05% trypsin-EDTA.

### 2.7 Cell viability assay

Cell viability was assessed using WST-1 as described previously (Ntsapi and Loos, 2021; Mthethwa *et al*., 2026). Cells (2 × 10^4^/well) attached for 24 hours in 96-well plates. After treatment, WST-1 (5 µL/well; Roche, CELLPRO-RO) was incubated for 2 hours at 37°C. Absorbance at 450 nm (600 nm reference) was measured using the GloMax Discover and blank-corrected. The following experiments used this assay, with untreated and solvent-matched vehicle controls.

#### 2.7.1 Basal cell viability

N2aWT cells received *H. erinaceus* extract (10-100 µg/mL) for 24 hours. Viability was normalised to the untreated control (100%).

#### 2.7.2 Aβ-induced injury model

Cells were exposed to Aβ_1-42_ (0.1, 0.5, 1, 2.5 or 5 µM) for 6, 12 or 18 hours. Viability was normalised to the untreated control (100%). The 6-hour exposure at the corresponding IC_50_ of 2.42 µM was selected for downstream experiments. Aβ presence in cell lysates was confirmed by Western blotting.

#### 2.7.3 EC_50_ determination for *H. erinaceus* and donepezil

After injury, Aβ-containing medium was replaced with medium containing *H. erinaceus* (10-100 µg/mL) or donepezil (6.25, 12.5, 25, 50 or 100 µM) for 24 hours. Responses were normalised to the Aβ-induced control (100%). EC_50_ estimates guided combination dose selection.

#### 2.7.4 Synergy analysis

Following injury, cells received checkerboard combinations for 24 hours at 0.125, 0.25, 0.5, 1 or 2 times each agent’s EC_50_. Zero-dose rows and columns supplied single-agent responses in a 6 × 6 matrix. Responses were normalised to the Aβ-induced control and evaluated as viability restoration using Bliss independence (Bliss, 1939), calculated in Excel and analysed in SynergyFinder Plus (Zheng *et al*., 2022).

### 2.8 Western blotting

Western blotting followed Ntsapi and Loos (2021) with the following treatment conditions. Cells (2 × 10^6^/T75 flask) attached for 24 hours and, after Aβ injury, received *H. erinaceus*, donepezil or the selected combination for 24 hours. Bafilomycin A1 (400 nM) was added to paired groups during the final 2 hours. RIPA lysates containing protease inhibitors were quantified by BCA assay and standardised to 1 mg/mL. Samples mixed 1:1 with 2× Laemmli buffer (20 µL/lane) were separated on 10% TGX gels and transferred to 0.2-µm nitrocellulose. Primary and HRP-conjugated secondary antibodies are listed in Supplementary Tables S1 and S2. ECL signals were captured with ChemiDoc and quantified using Image Lab 6.1 (Bio-Rad), normalised to β-actin.

### 2.9 Statistical analysis

Experiments were repeated independently three times; viability and BACE1 assays included technical triplicates. Data are mean ± SEM. IC_50_ and EC_50_ estimates were calculated for each independent experiment using variable-slope nonlinear regression in GraphPad Prism 10.6.1. Group differences were assessed by one-way ANOVA. Dunnett’s tests compared viability with the relevant control; Bonferroni tests compared each Western blot condition with its paired bafilomycin condition. Residual normality was assessed by Shapiro-Wilk testing. Significance was set at p < 0.05.

## 3. RESULTS

### 3.1 LC-MS profiling

Targeted LC-MS identified ergothioneine and trace hericenone D (Table 1; chromatograms in Supplementary Figure S1 and S2). Tentative untargeted annotations are listed in Supplementary Table S3.

**Table 1.** Targeted compounds in the methanolic *H. erinaceus* extract.

| Compound | Concentration (µg/mL) |
| --- | --- |
| Ergothioneine | 40.83 |
| Hericenone D | 0.0624 |

### 3.2 BACE1 inhibition and basal cell viability

The extract produced an increasing relative BACE1 inhibitory response with concentration. Basal N2aWT viability did not differ significantly from the untreated control after 24 hours (Figure 1).

**Figure 1.**
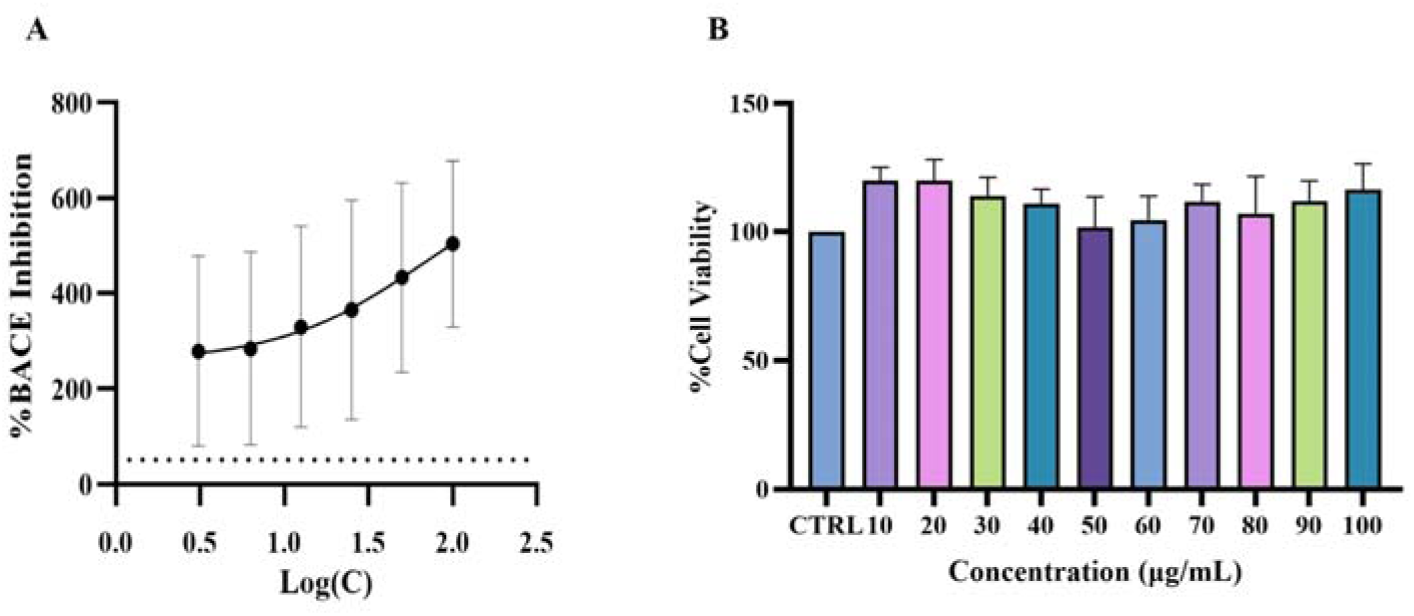
Effects of *H. erinaceus* on BACE1 activity and basal N2aWT viability. (A) FRET-based BACE1 response at 3.125-100 µg/mL, plotted against logarithmic extract concentration; IC_50_ was 69.97 ± 36.10 µg/mL. (B) WST-1 viability after 24 hours at 10-100 µg/mL, normalised to the untreated control (100%); no significant differences were observed. Data expressed as mean ± SEM (n = 3). Panel B: one-way ANOVA with Dunnett’s test against untreated control; p < 0.05.

### 3.3 Establishment of the Aβ-induced injury model

Aβ_1-42_ significantly reduced viability after 6 hours, with no significant reductions after 12 or 18 hours (Figure 2). The 6-hour IC_50_ of 2.42 µM was selected for subsequent experiments (Table 2). Aβ presence in cell lysates was confirmed by Western blotting (Supplementary Figure S3); IC_50_ curves are provided in Supplementary Figure S2.

**Figure 2.**
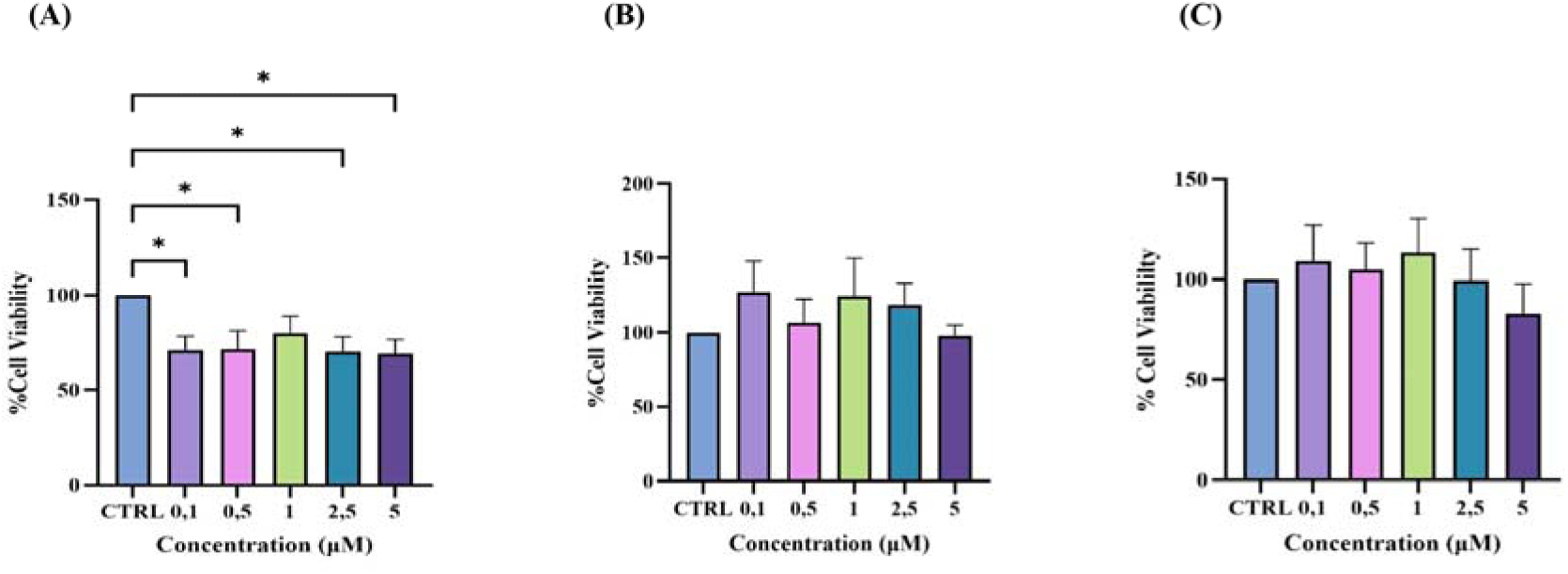
Effect of Aβ_1-42_ exposure on N2aWT cell viability. Cells were exposed to Aβ_1-42_ (0.1-5 µM) for (A) 6 hours, (B) 12 hours and (C) 18 hours. Cell viability was assessed using WST-1 and normalised to the untreated control (100%). At 6 hours, the overall effect was significant (*p = 0.0485), with significant reductions at 0.1, 0.5, 2.5 and 5 µM. No significant reduction was observed at 1 µM or after 12 or 18 hours. Statistical analysis used one-way ANOVA followed by Dunnett’s multiple comparisons test (*p < 0.05 vs. the untreated control). Data are presented as mean ± SEM from three independent experiments (n = 3). Analyses were performed using GraphPad Prism version 10.6.1.

**Table 2.**
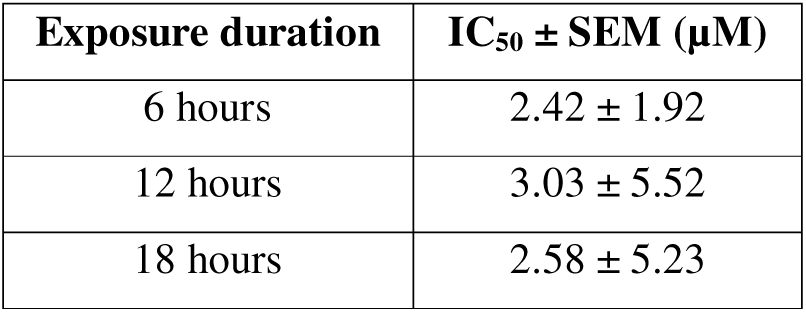
Aβ_1-42_ IC_50_ estimates at different exposure durations.

| Exposure duration | IC<sub>50</sub> ± SEM (μM) |
| --- | --- |
| 6 hours | 2.42 ± 1.92 |
| 12 hours | 3.03 ± 5.52 |
| 18 hours | 2.58 ± 5.23 |

### 3.4 EC_50_ determination for *H. erinaceus* and donepezil

Post-injury viability responses and EC_50_ estimates are shown in Figure 3.

**Figure 3.**
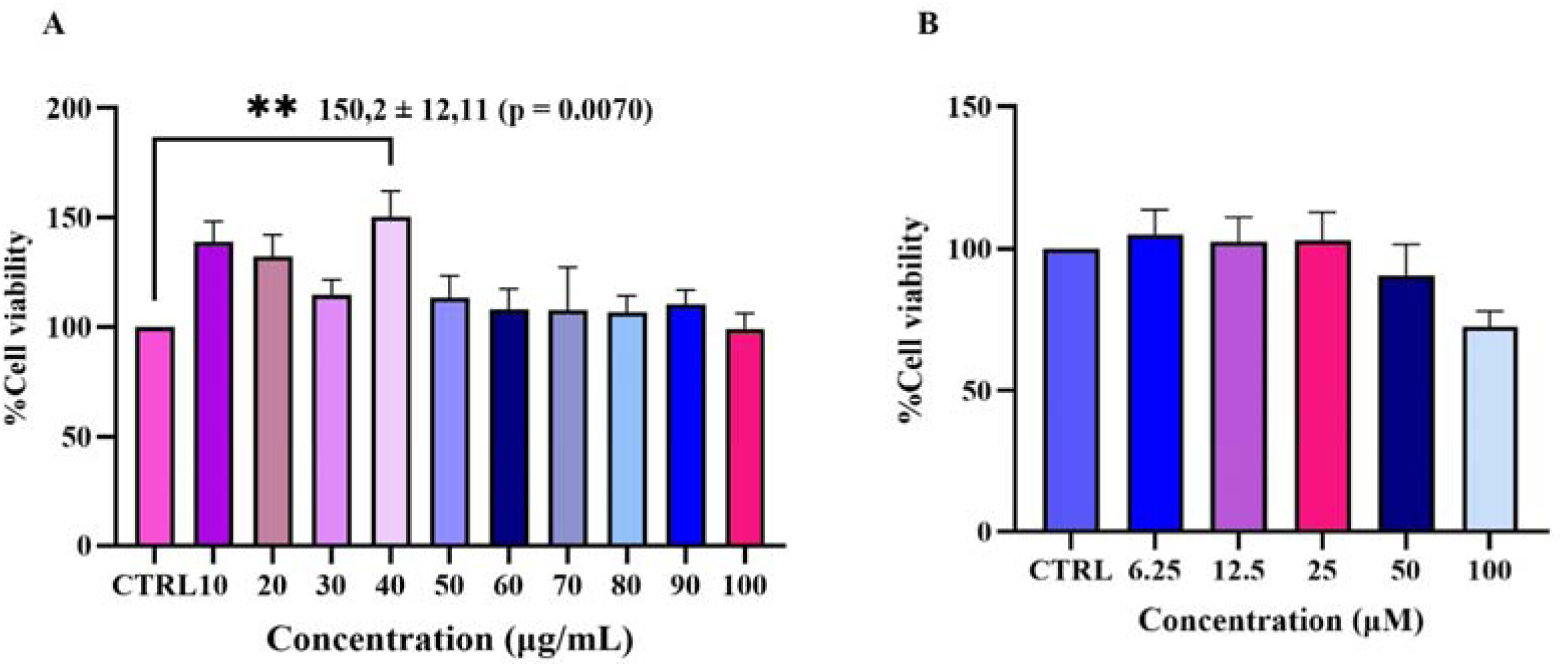
EC_50_ determination for *H. erinaceus* and donepezil. (A) Extract treatment (10-100 µg/mL) increased viability at 40 µg/mL (150.2 ± 12.11%; **p = 0.0070), with lower responses at higher concentrations; EC_50_ was 49.68 ± 5.01 µg/mL. (B) Donepezil (6.25-100 µM) produced responses near the Aβ-induced control at 6.25-25 µM and lower means at 50-100 µM; EC_50_ was 51.07 ± 5.07 µM. WST-1 responses were normalised to the Aβ-induced control (100%). Data are mean ± SEM (n = 3); one-way ANOVA with Dunnett’s test against the Aβ-induced control; p < 0.05.

**Figure 4.**
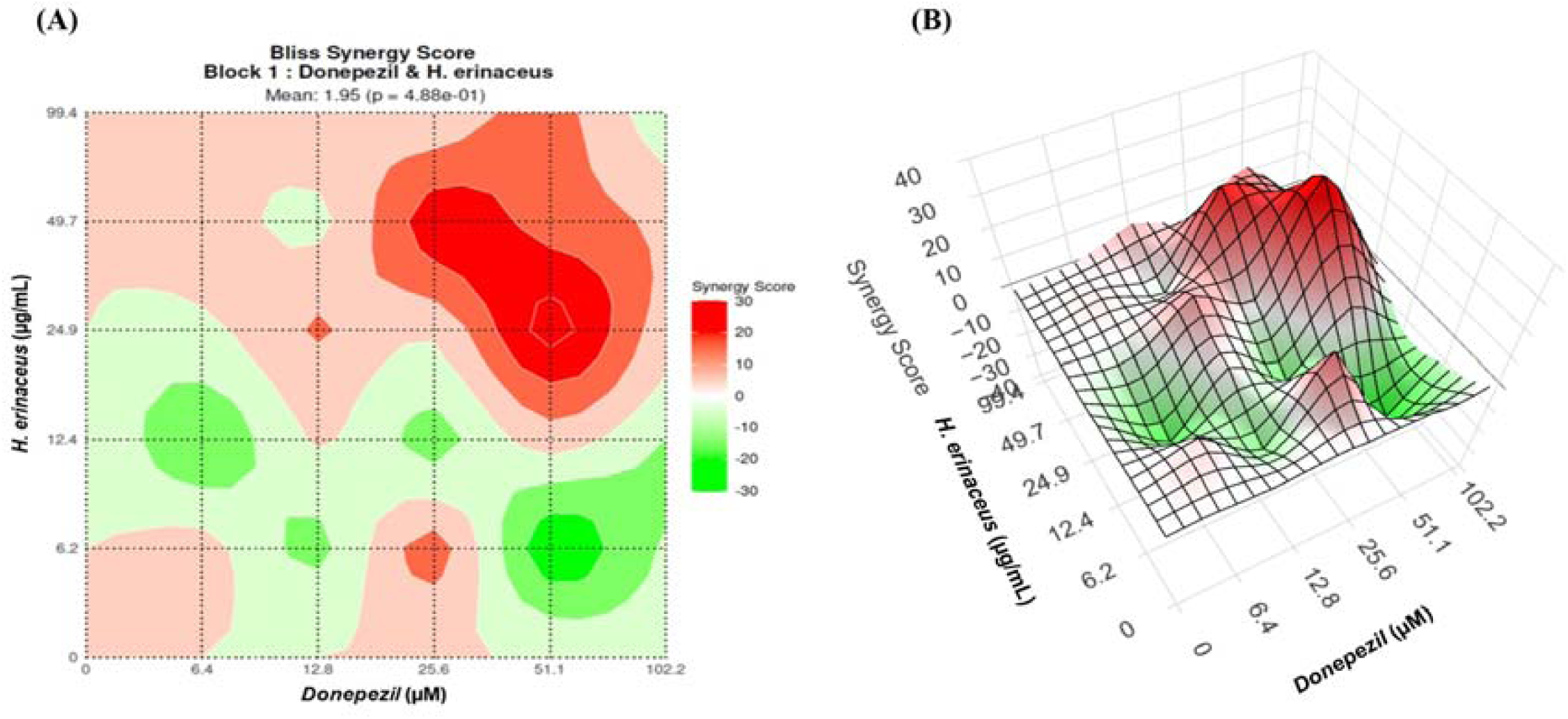
Bliss interaction maps for donepezil and *H. erinaceus*. (A) Two-dimensional heatmap and (B) three-dimensional surface. Red and green indicate positive and negative scores. The overall score was 1.95 (p = 0.488), within the additive range. Donepezil (51.1 µM) with extract (24.9 µg/mL) produced the highest local score (34.29) and viability and was selected for protein analysis. This 2:1 donepezil : extract ratio represents EC_50_ multiples of 1×:0.5×. Data are from three independent experiments. The 6 × 6 matrix and calculations are provided in Supplementary Table S4 and Supplementary Data S1.

### 3.5 Synergy analysis

Checkerboard viability responses were assessed using Bliss independence in SynergyFinder Plus to determine whether the agents interacted synergistically after injury.

### 3.6 Amyloid-associated and autophagy-related proteins

The selected pair was examined alongside each agent using Western blotting, with paired lysosomal blockade to investigate protein turnover. Extract and combination groups showed lower APP and Aβ means, with the lowest BACE1 mean in the combination (Figure 5). The combination also showed the most coordinated directional autophagy-related pattern, with comparatively higher ATG5 and LC3B, lower p62 and p62/Aβ accumulation with blockade (Figures 5 and 6). None of the tested paired comparisons was statistically significant.

**Figure 5.**
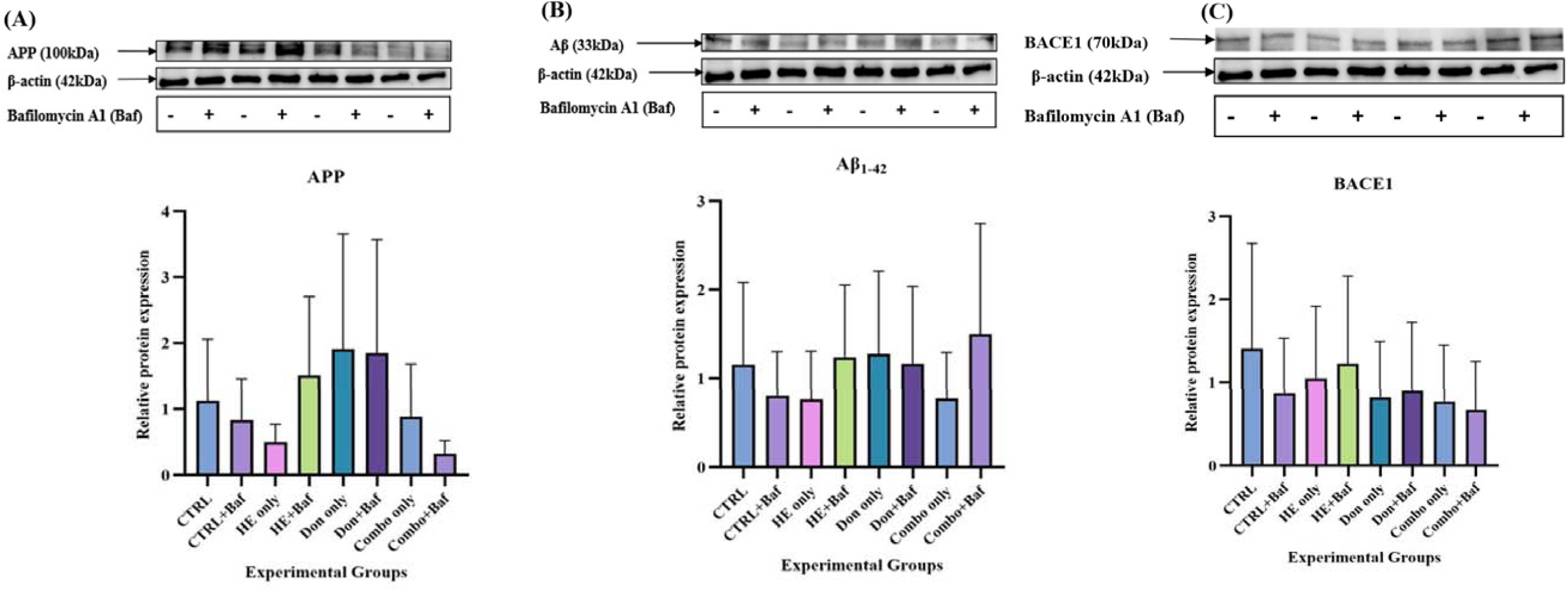
Amyloid-associated proteins after *H. erinaceus*, donepezil and combination treatment. Representative blots and densitometry for (A) APP, (B) Aβ_1-42_ and (C) BACE1, normalised to β-actin. Data are expressed as mean ± SEM (n = 3); one-way ANOVA with Bonferroni comparisons between each condition and its paired bafilomycin A1 condition; all p > 0.05. CTRL, Aβ-induced untreated control; HE, *H. erinaceus*; Don, donepezil; Combo, selected combination; Baf, bafilomycin A1.

**Figure 6.**
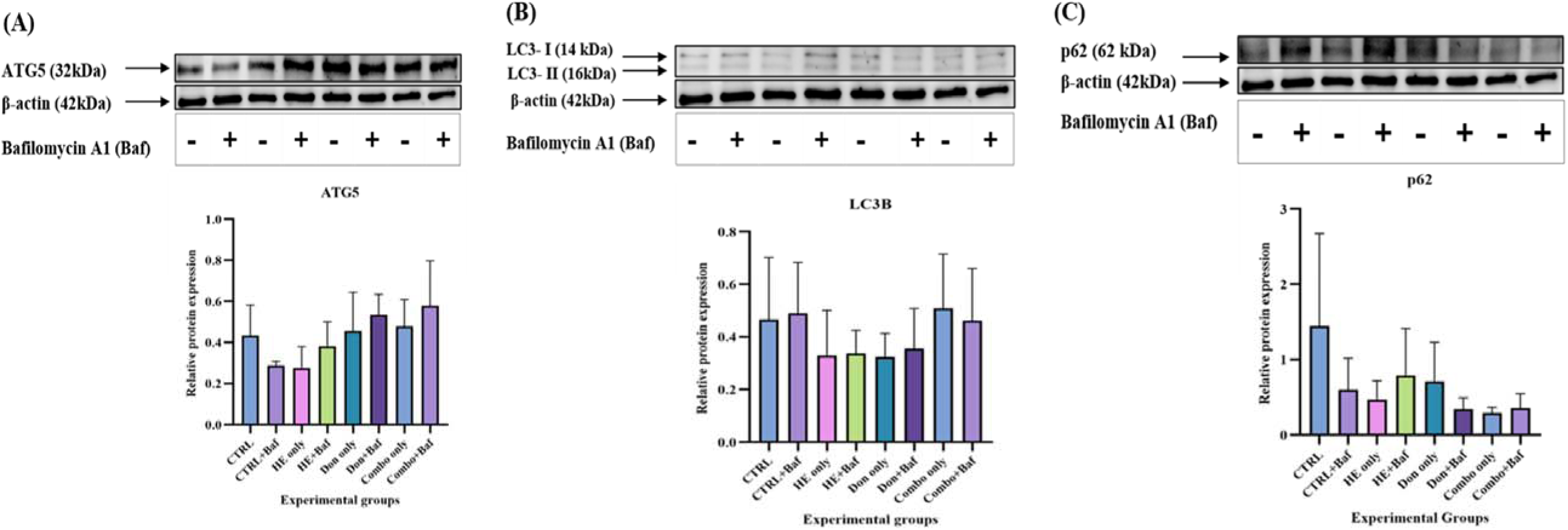
Autophagy-related proteins after *H. erinaceus*, donepezil and combination treatment. Representative blots and densitometry for (A) ATG5, (B) LC3B and (C) p62, normalised to β-actin. Data are expressed as mean ± SEM (n = 3); one-way ANOVA with Bonferroni comparisons between each condition and its paired bafilomycin A1 condition; all p > 0.05. CTRL, Aβ-induced untreated control; HE, *H. erinaceus*; Don, donepezil; Combo, selected combination; Baf, bafilomycin A1.

## 4. DISCUSSION

This study examined whether *H. erinaceus* could support recovery after amyloid-associated injury and complement donepezil through effects on production, clearance or both. The findings connect extract composition and apparent BACE1 inhibition with a responsive dose window, a concentration-specific combination effect and coordinated directional protein responses.

Ergothioneine and trace hericenone D provide chemical context for these responses. Untargeted annotations suggest broader chemical diversity but remain tentative without confirmatory MS/MS (Sumner *et al*., 2007); erinacine A was not analysed because a standard was unavailable. Ergothioneine reduced oxidative stress and plaque burden in 5xFAD mice (Whitmore *et al*., 2022) and improved Aβ clearance in the neuroretina (Wijesinghe *et al*., 2023), while selected hericenones stimulated NGF synthesis (Kawagishi *et al*., 1991). These studies support biological plausibility, but the crude extract’s activity cannot be assigned to one constituent.

The apparent concentration-dependent BACE1 response supports investigating reduced amyloidogenic processing, although its wide SEM and unstable fluorescence corrections limit precision. Natural-product constituents can interfere with fluorescence assays (Henrich and Beutler, 2013), warranting confirmation with an independent substrate-cleavage assay. Diling *et al*. (2017) identified predicted BACE1 interactions and reduced cellular BACE1 expression with isolated 3-hydroxyhericenone F in sodium azide-injured PC12 cells. This supports a BACE1-related action of *H. erinaceus* constituents, while the different compound and preparation prevent direct attribution of the present response.

The absence of significant basal viability reductions supported testing the extract after injury. Six-hour Aβ exposure provided measurable stress while retaining viability for subsequent treatment. Later exposures did not significantly reduce viability, potentially reflecting differences in peptide assembly and cellular responses. Vadukul *et al*. (2020) showed that Aβ conformation and assembly influence toxicity, illustrating why preparation and timing matter. The selected condition is therefore an acute amyloid-injury model for post-injury treatment assessment.

Following injury, the extract showed a biphasic response resembling its basal profile, with the strongest mean response at 40 µg/mL and less benefit at higher concentrations. This supports a responsive concentration window rather than continued benefit from dose escalation. Different constituents and stress responses may contribute, consistent with biphasic effects described in biological systems (Calabrese, 2013). The extract EC_50_ was numerically lower than its BACE1 IC_50_, but these values describe different endpoints. Cellular uptake, metabolism and responses to injury may contribute beyond enzyme inhibition, supporting examination of clearance alongside production.

Donepezil showed a narrower favourable profile, with declining mean viability at 50 and 100 µM. Noh *et al*. (2013) demonstrated protection during pretreatment through signalling beyond acetylcholinesterase inhibition. Differences in timing and cellular context may explain the more limited response here. Its EC_50_ guided combination dose selection.

Although the complete matrix was predominantly additive, the selected pair produced a greater-than-expected response and the highest observed viability. It retained donepezil at 1× EC_50_ while using 0.5× extract EC_50_, within the extract’s responsive range. This links combination treatment to dose optimisation: less extract was sufficient for the strongest response under these conditions. Inthachat *et al*. (2024) found target-dependent interactions between donepezil and *Diplazium esculentum* constituents, while Hong *et al*. (2022) reported benefits of a *Ginkgo biloba*-*H. erinaceus* formulation in a memory-impairment model. These comparisons support examining the selected combination’s cellular pathways.

Western blot patterns were interpreted descriptively because the tested comparisons between paired conditions with and without bafilomycin A1 were not significant. Lower APP and Aβ means with the extract and combination are compatible with reduced amyloid-associated protein accumulation. However, lower total APP may reflect altered synthesis or turnover and does not identify its cleavage route. Moreover, the measured Aβ pool may include retained introduced peptide and newly generated peptide. BACE1 modulation could limit new production while clearance would be needed to remove an existing pool; this distinction connects the enzyme findings to the post-injury model.

The combination’s lowest mean BACE1 abundance suggests a possible contribution from reduced enzyme availability, but protein abundance does not establish catalytic inhibition. Donepezil alone showed higher APP and Aβ means. Takada-Takatori *et al*. (2019) demonstrated that donepezil can alter APP endocytosis and favour non-amyloidogenic processing without reducing total APP. This illustrates why APP abundance and processing can diverge, although the present study did not measure cleavage products.

Lysosomal blockade helped examine protein turnover. Bafilomycin A1 disrupts acidification and autophagosome-lysosome fusion (Mauvezin and Neufeld, 2015), so its effects may involve trafficking and processing as well as degradation. The Aβ-induced control showed lower APP and Aβ means with blockade, whereas the extract showed directional accumulation. This difference is compatible with lysosome-dependent turnover contributing to the extract’s lower cargo abundance. Tzeng *et al*. (2016) also linked *H. erinaceus*-associated amyloid reduction to a clearance-related pathway through increased insulin-degrading enzyme, although that mechanism differs from autophagy.

ATG5 supports autophagosome formation, LC3-II associates with autophagosomal membranes, and p62 links cargo to the machinery and can itself be degraded. Their responses to blockade are more informative than abundance alone (Loos, du Toit and Hofmeyr, 2014). p62 also responds to stress-related synthesis through NRF2 (Jain *et al*., 2010), making coordinated interpretation necessary.

The extract’s lower p62 mean and directional accumulation with blockade aligned with its APP and Aβ responses. However, ATG5 was not clearly increased, and LC3-II was faint, leaving the formation component unresolved. Lin *et al*. (2023) reported SIRT1/ERK-associated autophagy and reduced protein accumulation with an alcoholic *H. erinaceus* mycelial preparation in A53T mice. These supports investigating an autophagy-related route, while the present extract pattern points more directly to bafilomycin-sensitive turnover.

The combination showed the most coordinated directional pattern: comparatively higher ATG5 and LC3B means, the lowest p62 mean, and accumulation of p62 and Aβ with blockade. APP and BACE1 remained comparatively low, suggesting that handling of existing peptide may accompany production-related changes. However, faint LC3-II and the absence of reliable separate LC3-I/LC3-II quantification prevent confirmation of increased flux.

The small biological sample size and variable protein signals limited precision. Independent viability measurements, greater replication, APP-cleavage analysis and LC3-II turnover measurements would strengthen interpretation. Separating introduced from newly generated Aβ would further resolve production and clearance.

## 5. CONCLUSION

This study suggests that methanolic *H. erinaceus* extract may attenuate amyloid-associated cellular changes through complementary effects on BACE1-associated amyloidogenic processing and intracellular clearance. In the Aβ_1-42_-induced N2a model, the extract supported post-injury viability recovery within a responsive concentration window. The selected combination with donepezil achieved the highest viability in the checkerboard matrix using a lower extract concentration and showed the most coordinated directional pattern of amyloid- and autophagy-related proteins.

Although the Western blot changes were not statistically significant, the aligned protein responses to lysosomal blockade were consistent with a possible clearance-related contribution. These integrated findings support further investigation of *H. erinaceus* as a potential adjunct to donepezil, with direct validation of amyloid processing and completed autophagic clearance needed to confirm the proposed dual response.

## 6. AVAILABILITY OF DATA AND MATERIALS

Data supporting the findings of this study are presented in the manuscript and supplementary material. Additional experimental data are available from the corresponding author upon reasonable request.

## 9. ACKNOWLEDGEMENTS

We thank the supervisory team for their guidance and support throughout the experimental work. We also thank Prof. Marieka Gryzenhout for providing the *H. erinaceus* material and Dr. Gabre Kemp for assistance with the LC-MS analysis.

## 10. LIST OF ABBREVIATIONS

### Abbreviation Definition

5xFAD: Mouse model expressing five familial Alzheimer’s disease mutations
AD: Alzheimer’s disease
AI: Artificial intelligence
ANOVA: Analysis of variance
APP: Amyloid precursor protein
APP/PS1: Amyloid precursor protein/presenilin 1 transgenic model
ATCC: American Type Culture Collection
ATG5: Autophagy-related protein 5
Aβ: Amyloid-β
Aβ_1-42_: Amyloid-β peptide comprising residues 1-42
BACE1: β-site amyloid precursor protein-cleaving enzyme 1
Baf: Bafilomycin A1
BCA: Bicinchoninic acid
Combo: Selected donepezil-*H. erinaceus* combination
CRediT: Contributor Roles Taxonomy
CTRL: Control
DMSO: Dimethyl sulfoxide
Don: Donepezil
EC□□: Half-maximal effective concentration
ECL: Enhanced chemiluminescence
EDTA: Ethylenediaminetetraacetic acid
ERK: Extracellular signal-regulated kinase
FRET: Fluorescence resonance energy transfer
HE: *Hericium erinaceus* extract
HPLC: High-performance liquid chromatography
HRP: Horseradish peroxidase
IC□□: Half-maximal inhibitory concentration
LC-MS: Liquid chromatography-mass spectrometry
LC3B: Microtubule-associated protein 1 light chain 3 beta
LC3-I: Non-lipidated form of LC3
LC3-II: Lipidated form of LC3
MS/MS: Tandem mass spectrometry
N2a: Neuro-2a mouse neuroblastoma cells
N2aWT: Wild-type Neuro-2a cells
NGF: Nerve growth factor
NRF2: Nuclear factor erythroid 2-related factor 2
p62: Sequestosome 1
PBS: Phosphate-buffered saline
PC12: Rat pheochromocytoma cell line
RIPA: Radioimmunoprecipitation assay buffer
SEM: Standard error of the mean
SIRT1: Sirtuin 1
TGX: Tris-Glycine eXtended
UFS: University of the Free State
WHO: World Health Organization
WST-1: Water-soluble tetrazolium salt 1

## 11. DECLARATION OF GENERATIVE AI AND AI-ASSISTED TECHNOLOGIES IN THE MANUSCRIPT PREPARATION PROCESS

During manuscript preparation, the authors used ChatGPT to assist with language editing, clarity and organisation. Generative AI was not used to generate experimental data or perform data analysis. The authors reviewed and edited the text and take full responsibility for the content of the manuscript.

## 12. CRediT AUTHOR STATEMENT

**Namatshego Moselakgomo:** Conceptualization, Writing - Original Draft, Software, Investigation. **Wenzile Sinothile Mthethwa:** Conceptualization, Investigation. **Claudia Ntsapi-Combrink:** Conceptualization, Supervision, Writing - Review & Editing, Funding acquisition. **Gabre Kemp:** Software, Methodology, Resources. **Marieka Gryzenhout:**

Writing - Review & Editing, Supervision, Visualization. **Sanet Van Zyl:** Writing - Review & Editing, Project administration, Resources

