## Supplementary Material for "The modulatory effects of *Hericium erinaceus* and donepezil on amyloid accumulation and autophagy-associated clearance in Aβ_1-42_-induced N2a cells"

**Supplementary Table S1: Primary antibodies used for Western blotting.**

| **Antibody name** | **Catalog number** | **Host species** | **Secondary antibody** | **Dilution factor** | **Band size (kDa)** |
| --- | --- | --- | --- | --- | --- |
| **LC3B Polyclonal Antibody** | PA1-46286 | Rabbit | Anti-Rabbit | 1:1000 | 14-16 |
| **ATG5 Polyclonal Antibody** | PA1-46178 | Rabbit | Anti-Rabbit | 1:1000 | 32 |
| **SQSTM1 (p62) Monoclonal Antibody** | GT1478 | Mouse | Anti-Mouse | 1:1000 | 62 |
| **Beta Amyloid Polyclonal Antibody (CT695) (APP)** | 51-2700 | Rabbit | Anti-Rabbit | 1:1000 | 100-140 |
| **BACE1 Polyclonal Antibody** | PA1-757 | Rabbit | Anti-Rabbit | 1:1000 | 70 |
| **Beta Actin Loading Control Monoclonal Antibody (BA3R)** | MA1-140 | Mouse | Anti-Mouse | 1:10000 | 42 |
| **Beta Amyloid (_1-42)_ Monoclonal Antibody** | AMY33 | Mouse | Anti-Mouse | 1:1000 | 33 |

| **Antibody name** | **Catalog number** | **Dilution factor** |
| --- | --- | --- |
| **Anti‐mouse IgG** | Cell Signalling # CST7076S | 1:1000 |
| **Anti‐rabbit IgG** | Cell Signalling # CST7074S | 1:1000 |

**Supplementary Table S2: The horseradish peroxidase-conjugated secondary antibodies used for Western blotting.**

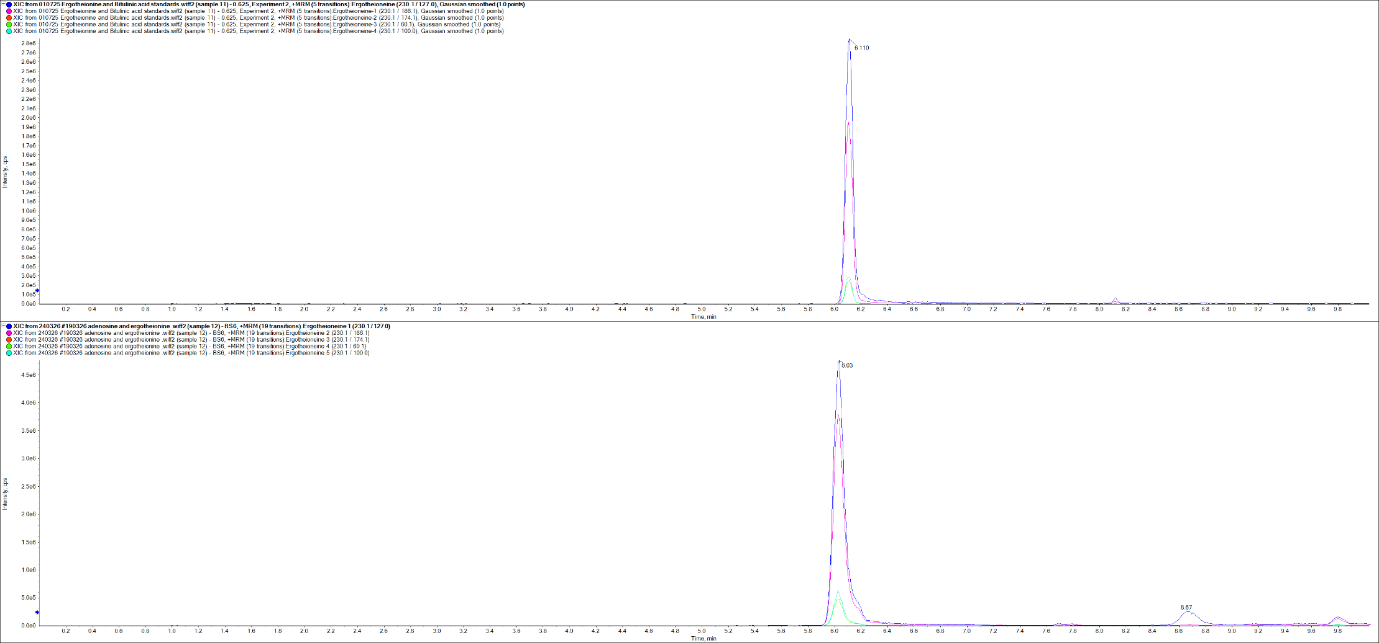

**A**

**B**

**Supplementary Figure S1: Targeted LC–MS/MS identification of ergothioneine in the *Hericium erinaceus* methanolic extract.** **(A)** MRM chromatogram of the ergothioneine reference standard, showing a peak at a retention time of 6.110 min. **(B)** MRM chromatogram of the methanolic *H. erinaceus* extract, showing the corresponding ergothioneine peak at 6.03 min. The overlaid traces represent five product-ion transitions from the precursor ion *m/z* 230.1 to *m/z* 127.0, 186.1, 174.1, 60.1 and 100.0. The comparable retention times and transition profiles supported the targeted identification of ergothioneine in the extract.

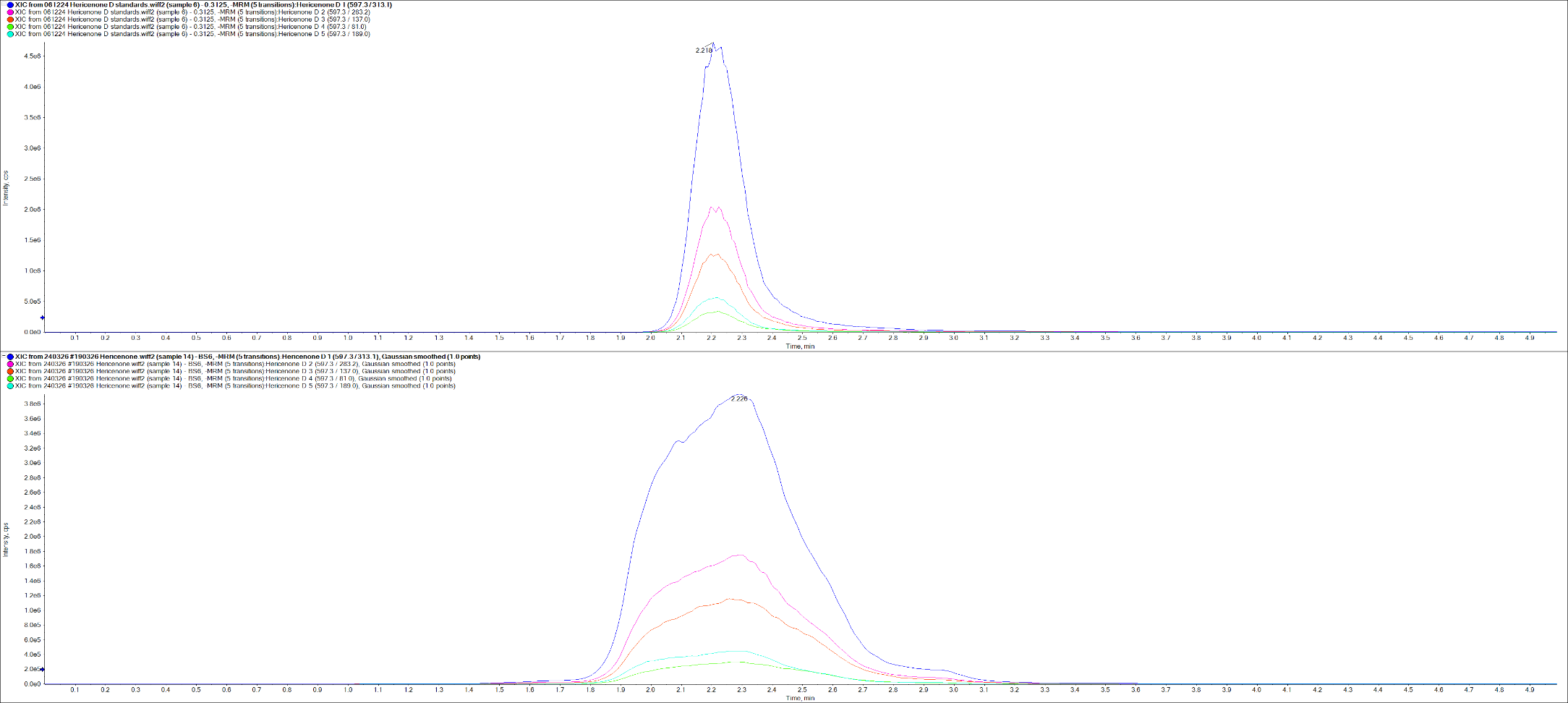

**B**

**A**

**Supplementary Figure S2: Targeted LC-MS/MS identification of hericenone D in the methanolic *Hericium erinaceus* extract.** **(A)** MRM chromatogram of the hericenone D reference standard, showing a peak at a retention time of 2.218 min. (B) MRM chromatogram of the methanolic *H. erinaceus* extract, showing the corresponding peak at 2.226 min. The overlaid traces represent five product-ion transitions from the precursor ion *m/z* 597.3 to *m/z* 313.1, 283.2, 137.0, 81.0 and 189.0. The comparable retention times and transition profiles supported the targeted identification of hericenone D in the extract.

**Supplementary Table S3: Untargeted positive-ion LC-MS/MS screening output.**

| **Replicate** | **Putative compound** | **Retention time (min)** | **Theoretical m/z** | **Observed m/z** | **Mass error (ppm)** |
| --- | --- | --- | --- | --- | --- |
| 1 | Aeruginascin | 14.247 | 300.1233456 | 300.1233561 | 0.035 |
| 1 | E-J | 20.154 | 467.2639446 | 467.2621509 | -3.839 |
| 1 | E-P | 20.097 | 493.2795947 | 493.2781587 | -2.911 |
| 1 | Lumichrome | 13.294 | 243.0876520 | 243.0876064 | -0.188 |
| 1 | Norharmane | 8.486 | 169.0760247 | 169.0763013 | 1.636 |
| 2 | Aeruginascin | 14.247 | 300.1233456 | 300.1235379 | 0.641 |
| 2 | E-J | 20.154 | 467.2639446 | 467.2626399 | -2.792 |
| 2 | E-P | 20.097 | 493.2795947 | 493.2778168 | -3.604 |
| 2 | Lumichrome | 13.294 | 243.0876520 | 243.0877411 | 0.366 |
| 2 | Norharmane | 8.486 | 169.0760247 | 169.0763850 | 2.131 |
| 3 | Aeruginascin | 14.247 | 300.1233456 | 300.1230060 | -1.132 |
| 3 | E-P | 20.097 | 493.2795947 | 493.2780655 | -3.100 |
| 3 | Lumichrome | 13.294 | 243.0876520 | 243.0874129 | -0.984 |
| 3 | Norharmane | 8.486 | 169.0760247 | 169.0763805 | 2.104 |

**Supplementary Table S4: Positive-ion library screening output.**

| **Replicate** | **Putative library match** | **Mass error (ppm)** | **Library score (%)** |
| --- | --- | --- | --- |
| 1 | Nicotinamide | -0.132 | 99.66 |
| 1 | Phenylalanine | -0.020 | 98.56 |
| 1 | Phenprobamate | -0.020 | 98.42 |
| 1 | Adenosine | -0.200 | 98.04 |
| 1 | Guanosine [Smart Confirmation] | 0.356 | 97.49 |
| 1 | Guanosine | 0.356 | 97.49 |
| 1 | Methyl linoleate | -0.668 | 95.02 |
| 1 | Glycerol trioleate (+NH3) | -0.098 | 94.53 |
| 1 | Isoleucine (L-isoleucine) | -0.729 | 91.00 |
| 1 | Adenine | -0.222 | 90.89 |
| 1 | Isoleucine | -0.729 | 87.31 |
| 1 | Leucine | -0.729 | 84.45 |
| 2 | Adenosine | 0.582 | 100.00 |
| 2 | Nicotinamide | -0.096 | 99.62 |
| 2 | Phenylalanine | 0.020 | 98.34 |
| 2 | Phenprobamate | 0.020 | 98.21 |
| 2 | Guanosine [Smart Confirmation] | 0.504 | 97.93 |
| 2 | Guanosine | 0.504 | 97.93 |
| 2 | Methyl linoleate | 1.144 | 92.86 |
| 2 | Adenine | 0.873 | 89.49 |
| 2 | Isoleucine (L-isoleucine) | 0.118 | 86.83 |
| 2 | Methyl linoleate | 0.366 | 83.56 |
| 2 | Cantharidin [Smart Confirmation] | 0.052 | 80.46 |
| 3 | Adenosine | -0.793 | 100.00 |
| 3 | Nicotinamide | -1.566 | 99.67 |
| 3 | Guanosine [Smart Confirmation] | 0.477 | 99.10 |
| 3 | Guanosine | 0.477 | 99.10 |
| 3 | Phenylalanine | -0.774 | 97.03 |
| 3 | Phenprobamate | -0.774 | 96.23 |
| 3 | Methyl linoleate | -0.953 | 93.16 |
| 3 | Isoleucine (L-isoleucine) | -0.716 | 92.77 |
| 3 | Isoleucine | -0.716 | 88.77 |
| 3 | Leucine | -0.716 | 86.74 |
| 3 | Adenine | 0.599 | 85.09 |
| 3 | Muscone | 0.333 | 84.77 |
| 3 | Nicotinic acid | -0.599 | 81.86 |

**Supplementary Table S5: Untargeted negative-ion LC-MS/MS screening output.**

| **Replicate** | **Putative compound** | **Retention time (min)** | **Mass error (ppm)** |
| --- | --- | --- | --- |
| 1 | H-A | 17.838 | 0.692 |
| 1 | H-A | 24.834 | 1.412 |
| 1 | H-B | 19.516 | -1.216 |
| 1 | H-B | 23.960 | 1.226 |
| 1 | H-C/F | 23.982 | -0.790 |
| 1 | H-E/H | 26.444 | 0.057 |
| 1 | H-G/D | 23.371 | 0.403 |
| 1 | H-G/D | 23.986 | 0.121 |
| 1 | H-G/D | 26.853 | 0.835 |
| 1 | H-I | 23.143 | 0.099 |
| 1 | H-I | 24.016 | 0.944 |
| 1 | H-I | 25.862 | -0.417 |
| 1 | H-J | 21.081 | 0.237 |
| 2 | H-A | 17.838 | -0.077 |
| 2 | H-A | 24.834 | 0.688 |
| 2 | H-B | 19.516 | -3.079 |
| 2 | H-B | 23.960 | 0.816 |
| 2 | H-C/F | 23.982 | -0.681 |
| 2 | H-E/H | 26.444 | -0.515 |
| 2 | H-G/D | 26.853 | 0.379 |
| 2 | H-I | 23.143 | -0.535 |
| 2 | H-I | 24.016 | 0.648 |
| 2 | H-J | 21.081 | 0.853 |
| 3 | H-A | 17.838 | 0.961 |
| 3 | H-A | 24.834 | 1.200 |
| 3 | H-B | 19.516 | 0.016 |
| 3 | H-B | 23.960 | 1.857 |
| 3 | H-E/H | 26.444 | 0.523 |
| 3 | H-I | 25.862 | -0.168 |
| 3 | H-J | 21.081 | 0.800 |
| 3 | Lumichrome | 8.313 | -3.825 |

**Supplementary Table S6: Negative-ion library screening output.**

| **Replicate** | **Putative library match** | **Mass error (ppm)** | **Library score (%)** |
| --- | --- | --- | --- |
| 1 | Adenosine | 0.123 | 100.00 |
| 1 | Guanosine | -0.651 | 100.00 |
| 1 | Adenine | -0.895 | 99.91 |
| 1 | L-Tryptophan | -1.297 | 99.09 |
| 1 | Adenine | -0.782 | 96.04 |
| 1 | D-Sorbitol | -3.628 | 94.63 |
| 1 | Xylitol | -2.753 | 94.07 |
| 1 | L-Malic acid | -5.033 | 93.14 |
| 1 | Mannitol | -3.628 | 93.03 |
| 1 | Rhein [Smart Confirmation] | -0.052 | 91.95 |
| 1 | 2-Hydroxyadenosine | -0.651 | 88.33 |
| 2 | Adenosine | -0.465 | 100.00 |
| 2 | Adenine | 0.814 | 99.17 |
| 2 | Guanosine | -0.706 | 98.64 |
| 2 | L-Malic acid | -2.006 | 97.75 |
| 2 | L-Tryptophan | -0.947 | 97.34 |
| 2 | Rhein [Smart Confirmation] | -0.351 | 95.12 |
| 2 | Xylitol | -0.768 | 93.76 |
| 2 | Isoleucine [Smart Confirmation] | -0.450 | 93.41 |
| 2 | Isoleucine [Smart Confirmation] | -0.450 | 93.41 |
| 2 | Isoleucine [Smart Confirmation] | -0.450 | 93.41 |
| 2 | Isoleucine [Smart Confirmation] | -0.450 | 93.41 |
| 2 | Isoleucine | -0.450 | 93.41 |
| 2 | Isoleucine | -0.450 | 93.41 |
| 2 | Adenine | -0.737 | 92.40 |
| 2 | Mannitol | -2.115 | 91.83 |
| 2 | D-Sorbitol | -2.115 | 90.28 |
| 2 | Rhein [Smart Confirmation] | 2.035 | 89.74 |
| 2 | 2-Hydroxyadenosine | -0.706 | 80.39 |
| 3 | Adenosine | 0.628 | 100.00 |
| 3 | L-Tryptophan | 0.733 | 100.00 |
| 3 | Guanosine | 0.339 | 100.00 |
| 3 | Adenine | 0.795 | 99.52 |
| 3 | L-Malic acid | -2.348 | 96.81 |
| 3 | Adenine | -0.348 | 94.22 |
| 3 | Xylitol | -0.765 | 94.15 |
| 3 | Rhein [Smart Confirmation] | 1.554 | 91.70 |
| 3 | D-Sorbitol | -1.216 | 91.06 |
| 3 | Mannitol | -1.216 | 88.28 |
| 3 | Uridine | 0.798 | 84.82 |
| 3 | Rhein [Smart Confirmation] | 2.732 | 84.45 |

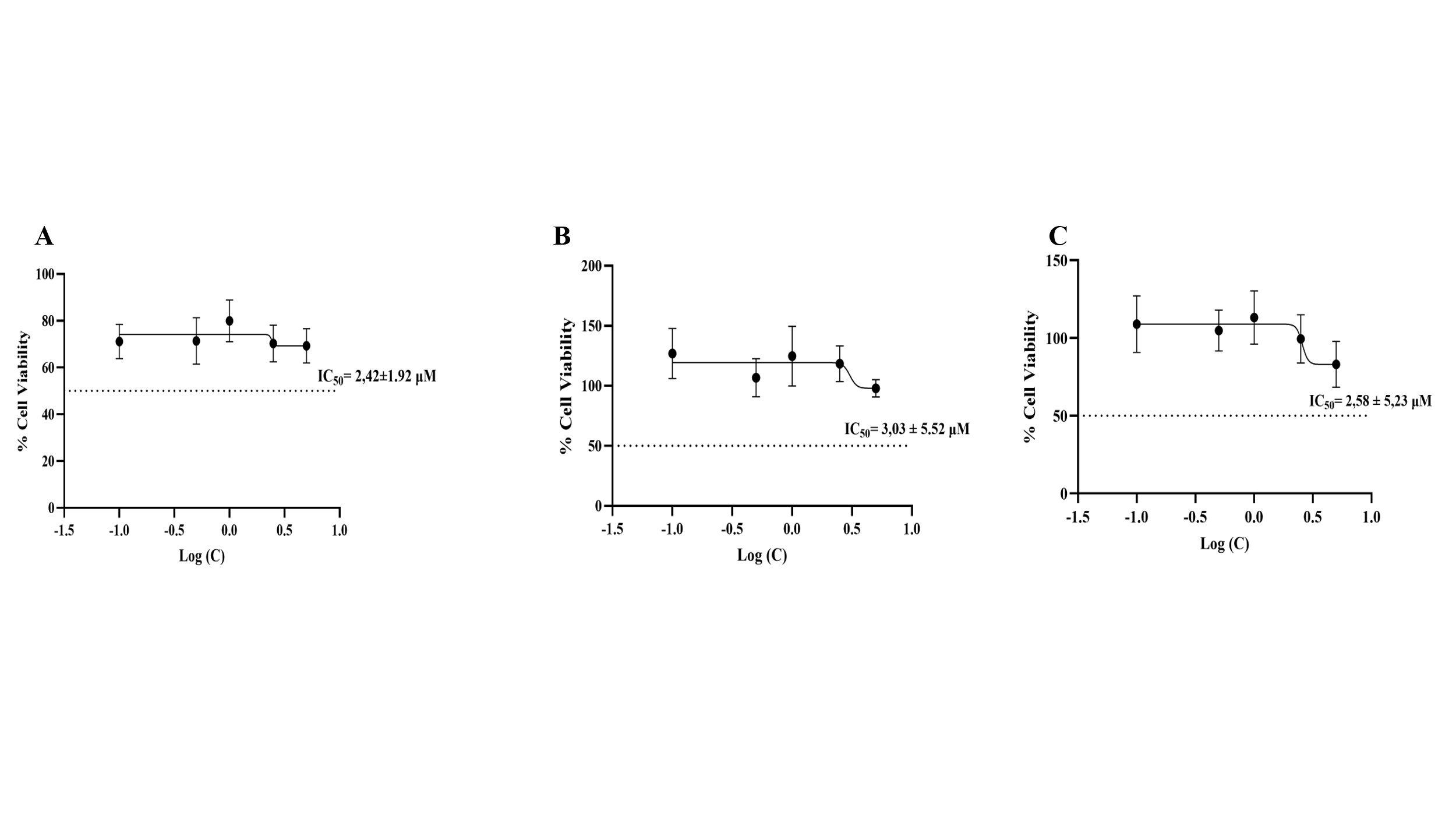

**Supplementary Figure S3: Nonlinear-regression curves for N2aWT cell viability following Aβ_1-42_ exposure.** N2aWT cells were exposed to increasing concentrations of Aβ_1-42_ (0.1-5 µM) for (A) 6, (B) 12 and (C) 18 hours. Cell viability was determined using the WST-1 assay and expressed relative to the control. Nonlinear-regression curves were fitted to the complete concentration-response datasets to estimate the IC_50_ at each exposure interval. Data are presented as mean ± SEM (n = 3).
